# First description of the sexual stage of *Venturia effusa*, causal agent of pecan scab

**DOI:** 10.1101/785790

**Authors:** Nikki D. Charlton, Mihwa Yi, Clive H. Bock, Minling Zhang, Carolyn A. Young

**Author notes:** Carolyn A. Young (corresponding author).

## Abstract

*Venturia effusa*, cause of pecan scab, is the most prevalent disease of pecan in the southeastern USA; epidemics of the disease regularly result in economic losses to the pecan industry. Recent characterization of the mating type distribution revealed the frequency of the *MAT* idiomorphs are in equilibrium at various spatial scales, indicative of regular sexual recombination. However, the occurrence of the sexual stage of *V. effusa* has never been observed, and the pathogen was previously believed to rely entirely on asexual reproduction. To explore the existence of a sexual cycle, we paired opposite mating types on oatmeal culture media. In initial experiments, cultures were incubated at 24 C for 2 mo for hyphal interactions to occur between mating types and then maintained at 4 C for 4 mo. Immature pseudothecia were initially observed but following exposure to a 12 h photoperiod for 2 weeks at 24 C, asci and ascospores developed. Further experiments explored the effect of time on pseudothecial development with 4 mo at 4 C as the optimal requirement. The results of this study demonstrate the heterothallic nature of *V. effusa.* Following experiments investigated progeny from a sexual cross of an albino and a wild-type isolate. Evaluation of isolate pigmentation, mating type, and multilocus genotyping of single ascospore progeny provided evidence that recombination occurred within the sexual crosses. The impact of determining the source of the overwintering ascostroma will aid in management decisions to reduce the primary inoculum in the disease cycle.

## INTRODUCTION

Pecan scab, caused by *Venturia effusa* (G. Winter) Rossman & WC Allen (previously *Fusicladium effusum* G. Winter) (Rossman et al. 2016), is the most important biotic, yield-limiting issue impacting pecan production in the southeastern United States. Pecan is one of the few plants native to North America that has developed into a significant and lucrative specialty crop (Wood et al. 1990). Pecan trees can live and bear edible fruit for over a hundred years, if well managed (Wells 2017). The value of the pecan crop in the US in 2017 was estimated at $ 709,218,000 (NASS, 2017). The estimated impact of scab can be measured in two ways – firstly the direct impact of the disease reducing yield, and secondly the cost of fungicides to manage it. For example, in an average scab year like 2014, statewide production in GA was valued at $ 313,313,250, the yield loss to the disease was estimated at $ 3.1 million, and the cost of fungicides for control was estimated at $ 22.6 million (Brock and Brenneman 2014). In severe epidemic years both costs are elevated (Bock et al. 2017a).

*V. effusa* was first described in 1885 on mockernut in Illinois (Winter 1885). The pathogen has been considered to perpetuate purely through asexual reproduction by repeated production of conidia throughout its life cycle. The conidia are spread by wind and splash dispersal and can infect under humid, moist conditions; they are a key component of the polycyclic nature of this disease (Gottwald and Bertrand 1983; Gottwald 1985). The asexual lifecycle can be completed in as little as 7 to 10 days, and epidemics develop as inoculum builds up throughout the season under favorable environmental conditions (sporulation, dispersal, germination, infection and lesion development) (Gottwald 1982; Gottwald and Bertrand 1982). When severe, the epidemics may affect the number and quality of harvested nuts (Gottwald and Bertrand 1983; Stevenson and Bertrand 2001).

The population genetic structure and diversity of a pathogen provides valuable information and insight into the epidemiology of a plant disease. Asexual and sexual reproduction influence the population structure, and ultimately its ability to adapt to different selection (McDonald and Linde 2002; McDonald and Mundt 2016). Understanding the spatial distribution and occurrence of clones provides insight into the modes of reproduction of a pathogen (McDonald and McDermott 1993; Milgroom 1996; Bock et al. 2016a). Sexual reproduction introduces recombination and diversity into a population. Pathogens with mixed reproduction systems (asexual and sexual) are proposed to have the highest risk of evolution allowing the introduction of new genotypes through recombination and distribution of these genotypes through clonal propagation (McDonald and Linde 2002). These high-risk pathogens are considered capable of evading major resistance genes or evolving rapidly to overcome control methods such as specific fungicide types.

Recent sequencing efforts (Bock et al. 2016b) have been used to facilitate insight into the population genetic structure and dynamics of *V. effusa* in the southeastern USA. Evaluation of hierarchically collected isolates from multiple orchards using microsatellite markers has shown substantial and evenly distributed genetic diversity among orchard populations (Bock et al. 2017b). Such distribution and nature of the genetic diversity are characteristic of a sexually reproducing species (McDonald et al. 1995; Linde et al. 2002; Gladieux et al. 2008; Gladieux et al. 2010). In addition, *V. effusa* is proposed to have a heterothallic mating system as isolates were observed to contain either the *MAT1-1-1* or *MAT1-2-1* mating type idiomorphs, which also were found to be in equilibrium within the orchards examined (Bock et al. 2018; Young et al. 2018). Based on these observations we propose that *V. effusa* must have a functional sexual stage that is yet to be observed in the field. Closely related species of *Venturia* (Schubert et al. 2003), including *V. inaequalis* on apple, *V. nashicola* on pear, and *V. carpohila* on peach and apricot have a characterized sexual stages in the field in at least some areas (Fisher 1961; Umemoto 1990; MacHardy 1996; González-Domínguez et al. 2017). In some areas, ascospores of *V. nashicola* may play a bigger role in epidemics of pear scab than previously thought (Lian et al. 2006; Lian et al. 2007). In the case of *V. inaequalis*, the sexual stage is often the primary source for the epidemic (MacHardy 1996), although conidia may also contribute (Passey et al. 2017). Control measures for apple scab in some areas may be aimed at the sexual stage to reduce impact of the disease on apple (Holb 2006).

Current management practices for scab on pecan rely on the use of fungicides (Bock et al. 2017a). Fungicide resistance is already documented for *V. effusa* to benomyl (Littrell 1976) and the DMIs (Reynolds et al. 1997). More recent work indicates that there is reduced sensitivity to tin-based products (Standish et al. 2018), and a risk of resistance development to QoIs through specific mutations other than the more common G143A substitution may exist (Standish et al. 2016). Thus, there is reason for serious concern that fungicide resistance may become an increasingly greater issue based on what we currently know about the genetic diversity of populations of *V. effusa*. In order to develop effective disease management strategies for pecan scab, the disease cycle must be well understood. The genetic diversity (Bock et al. 2017b) and the equilibrium of the mating type idiomorphs (Bock et al. 2018; Young et al. 2018) suggests sexual reproduction occurs in populations of *V. effusa* across the southeastern United States; therefore we postulate a role for ascospores in the epidemiology of *V. effusa*. Knowing the timing of production and infection process of ascospores, and their role in the epidemiology of pecan scab may prove decisive in aiding management of this devastating disease.

In this study we present *in vitro* evidence of the sexual stage of *V. effusa*. We describe the methodology to produce the sexual stage and utilize microscopy to demonstrate the developmental stages of the pseudothecium. In addition, single ascospore progeny were selected and screened using microsatellite markers to demonstrate viability and evidence of recombination through sexual reproduction.

## MATERIALS AND METHODS

### Isolates

Sixteen isolates of *V. effusa* were used in this study (TABLE 1). Seven isolates were previously characterized in a population study (Bock et al. 2017b). One albino isolate (FRT5LL7-Albino, a natural variant selected as an albino sector from wild-type, melanized isolate FRT5LL7) was isolated from a commercial orchard of the pecan cultivar Desirable near southwest Albany, GA, and the remaining eight isolates were sampled from scab-diseased leaves or stems of pecan from orchards in Oklahoma and Texas. Methods for isolation have been previously described in detail (Bock et al. 2017b). Briefly, monoconidial isolates were obtained from individual lesions by spreading a conidial suspension on water agar (WA) amended with antibiotics (lactic acid [0.50 mL/L], streptomycin [0.20 g/L], tetracycline [0.05 g/L] and chloramphenicol [0.05 g/liter]), and incubating for 1 day at 25 C. Under a microscope (50×), a single germinating conidium was carefully removed and transferred to antibiotic amended (as for WA) potato dextrose agar (PDA). Isolates were maintained on both PDA and oatmeal agar (OMA; 30 g Gerber’s organic baby oatmeal and 20 g agar per liter) at 25 C with a 12 h photoperiod; suspensions were prepared for long term storage on silica at −20 C.

**Table 1.**
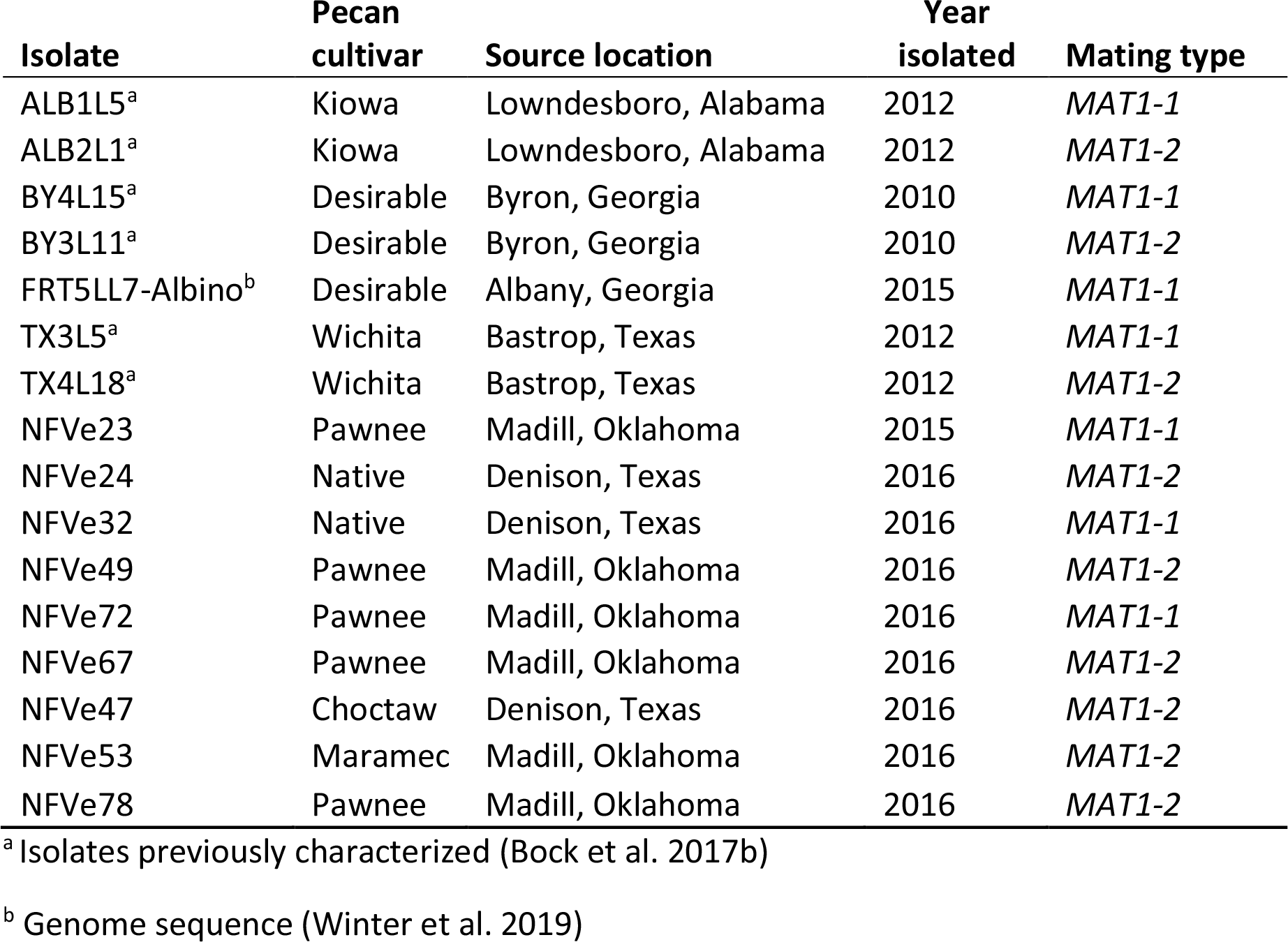
*Venturia effusa* isolates used in this study

### DNA extraction and PCR

Mycelium was scraped from a 2-3 wk old colony of *V. effusa* growing on PDA and DNA was extracted using the *Quick*-DNA Fungal/Bacterial Miniprep Kit (Zymo Research, Irvine, CA) according to manufacturer’s instructions. Primers for mating type (*MAT1-1* and *MAT1-2*) and the housekeeping gene *TUB2* were multiplexed as previously described (Young et al. 2018) to identify the mating type for each isolate.

### Mating assay

Isolates of opposite mating type, self and non-self-same mating type were paired on OMA and PDA on 9 cm quadrant petri plates (TABLE 2). Preliminary pairings were established by placing adjacent plugs of agar with mycelia of each mating type on OMA (Fig. 1). Once the sexual stage had been observed, the preliminary methodology was improved for future pairings using ground inoculum. The ground inoculum was prepared by scraping mycelium from a 2-3 wk old colony of *V. effusa* grown on PDA and placed in a 2 ml tube with three 3 mm glass beads and 500 μL of sterile potato dextrose broth. The samples were subject to grinding using a Qiagen TissueLyser (QIAGEN Inc., Valencia, CA) at 30 Hz for 6-9 min. Three inoculation points for each isolate per quadrant were made by pipetting 2 μL of inoculum onto the OMA (Fig. 1). The preliminary pairings using plugs of agar were placed in a 24 C incubator with a 12 h photoperiod for 4-8 wk. When using the ground-mycelium drop inoculation methodology, the pairings were placed in the 24 C incubator for 2 wk. Once the paired isolates had sufficiently grown for hyphal overlap and interactions, the plates were wrapped in aluminum foil and placed at 4 C for 4 mo. After the 4 mo cold treatment, the interaction zone between the paired isolates was checked for any evidence of sexual fruiting structures, and the plates were placed in an incubator at 24 C with a 12 h photoperiod. The plates were checked for evidence of sexual fruiting structure every few days. Pseudothecia were observed and randomly selected from the zone between opposite mating types under a dissecting microscope. The pseudothecia were placed in a drop of sterile water on a glass slide. A coverslip was placed over the drop and the pseudothecia observed with a Leica CME microscope at 40× magnification. Slight pressure was applied using a cotton swab to split the pseudothecium to evaluate the maturity of the ascocarp (immature = no asci, partially mature = contents contain some asci with and without ascospores, fully mature = contents contain many asci with ascospores). A graphic timeline of the experimental conditions required for *in vitro* sexual reproduction in *V. effusa* is presented (Fig. 1).

**Table 2.** Crosses of *Venturia effusa*

| **Isolate** | **Mating type** | **Isolate** | **Mating type** | **No. of replicated crosses** |
| --- | --- | --- | --- | --- |
| ALB1L5<sup>a</sup> | *MAT1-1* | ALB2L1<sup>a</sup> | *MAT1-2* | 5 |
| ALB1L5<sup>a</sup> | *MAT1-1* | BY3L11<sup>a</sup> | *MAT1-2* | 5 |
| BY4L15<sup>a</sup> | *MAT1-1* | BY3L11<sup>a</sup> | *MAT1-2* | 5 |
| TX3L5<sup>a</sup> | *MAT1-1* | TX4L18<sup>a</sup> | *MAT1-2* | 3 |
| NFVe23 | *MAT1-1* | NFVe24 | *MAT1-2* | 1 |
| NFVe23 | *MAT1-1* | NFVe49 | *MAT1-2* | 1 |
| NFVe32 | *MAT1-1* | NFVe24 | *MAT1-2* | 1 |
| NFVe72 | *MAT1-1* | NFVe67 | *MAT1-2* | 1 |
| Albino | *MAT1-1* | NFVe47 | *MAT1-2* | 1 |
| Albino | *MAT1-1* | NFVe53 | *MAT1-2* | 1 |
| Albino | *MAT1-1* | NFVe78 | *MAT1-2* | 1 |
| ALB1L5<sup>b</sup> | *MAT1-1* | BY4L15<sup>b</sup> | *MAT1-1* | 2 |
| ALB2L1<sup>b</sup> | *MAT1-2* | BY3L11<sup>b</sup> | *MAT1-2* | 2 |

<sup>a</sup>Preliminary study<sup>b</sup>Non-self-same mating types were paired as controls

**FIGURE 1.**
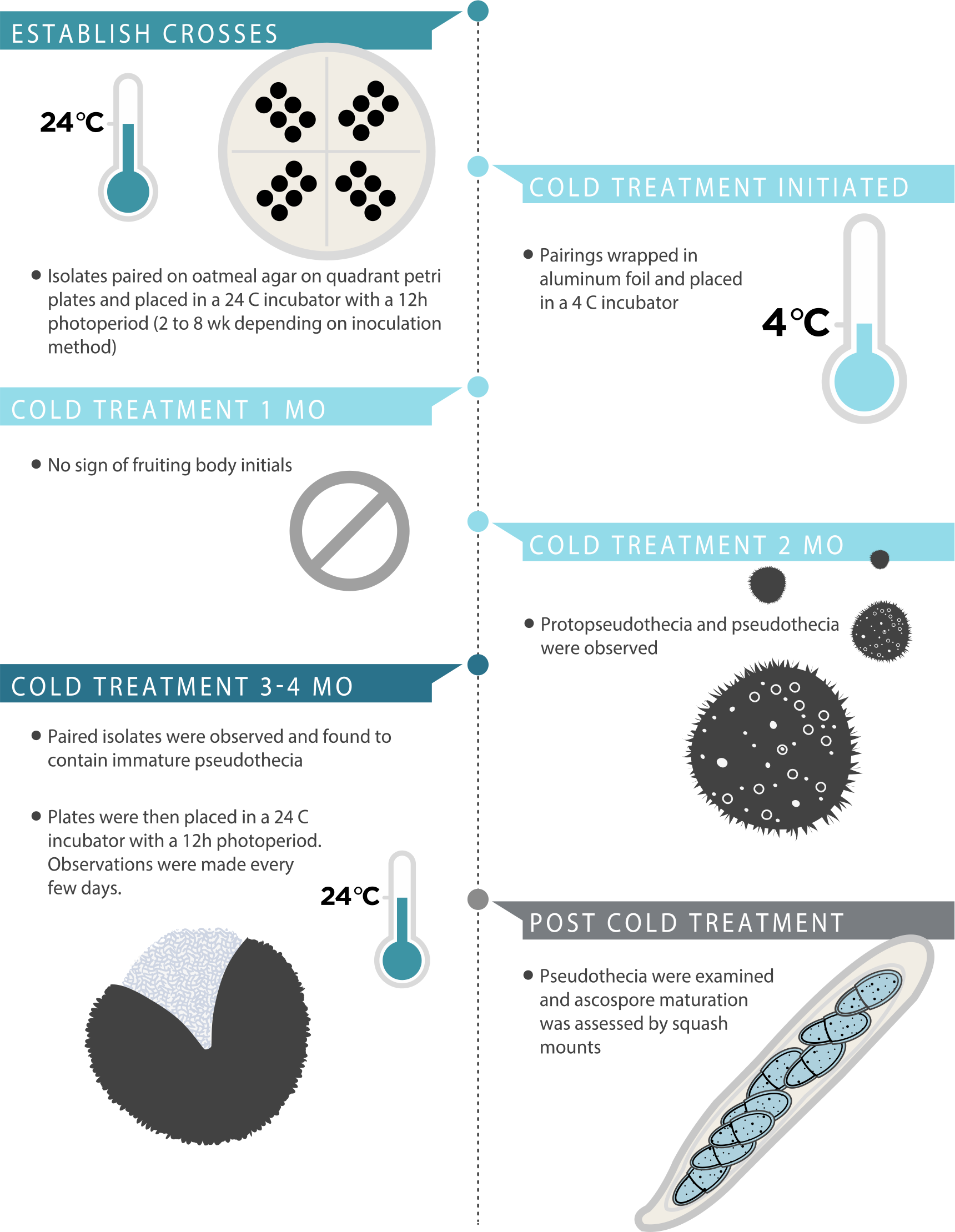
Graphic representation depicting the time and temperature required for *in vitro* sexual reproduction in *Venturia effusa*.

### Effect of time on pseudothecia development

Pairings were repeated to observe the effect of the length of cold treatment. Pairings were exposed to either 1, 2, 3 or 4 mo cold treatment at 4 C, before transfer to an incubator at 24 C with a 12 h photoperiod for 2 wk when they were observed for pseudothecial development. Pseudothecia were examined and ascospore maturation was assessed by squash mounts. Pseudothecia were rated as immature = no asci, partially mature = a few asci with ascospores and fully mature = numerous asci with ascospores.

### Sectioning for microscopy

Agar blocks (ca. 5 mm × 7 mm) containing pseudothecia were taken and frozen in liquid nitrogen. The block was mounted and embedded in Tissue-Tek^®^ O.C.T. Compound (Electron Microscopy Sciences, Hatfield, Pennsylvania). Twenty micrometer sections were cut on a Leica CM1850 Cryostat (Leica Biosystems Inc., Buffalo Grove, IL) and mounted on microscopy slides. The sections were stored at −20 C until processing when they were stained with toluidine blue. The cross and longitudinal sections were observed under an Olympus BX41 microscope using LUC Plan FL N 40x/0.60 NA and Plan N 100x/1.25 NA (oil) objectives. Images were captured using an Olympus DP71 CCD camera with DP Controller software (Olympus America Inc.). Subsequently, selected sections were observed using Zeiss Axio Imager M2 upright microscopes (Carl Zeiss, Germany) with Plan-Neofluar 40x/0.75 NA and Plan-Neofluar 100X/1.30 NA (oil) objectives equipped with DIC optics. Images were captured using an Axiocam 503 mono camera with ZEN 2 (Blue edition) Image acquisition and processing software.

### Isolation of ascospore progeny

Three crosses were chosen for ascospore isolation (NFVe23 × NFVe24, FRT5LL7-Albino × NFVe47, and FRT5LL7-Albino × NFVe78). Individual pseudothecia were squashed and the contents were plated on 2% water agar. Single germinating ascospores were arbitrarily isolated using a microdissection needle and transferred to PDA and grown at 24 C with a 12h photoperiod. Ascospore progeny were stored on silica at −20 C.

### DNA extraction and multilocus genotyping of ascospore progeny

Isolates were grown on PDA for 2-3 wk. DNA was extracted from freeze dried fungal mycelia using the Qiagen MagAttract 96 DNA Plant Core Kit (QIAGEN Inc., Valencia, CA) according to the manufacturer’s protocol. Parental and single ascospore progeny were screened for polymorphisms using 32 microsatellite markers as described previously (Bock et al. 2016a; Bock et al. 2017b). PCR was performed in a 10 μL volume containing template DNA (approximately 2 ng), 0.5 U GoTaq™ DNA Polymerase (Promega Corp., Madison, Wisconsin), 1× Colorless GoTaq™ PCR Buffer, 150 nM of each dNTP (Promega Corp.), 100 nM of the reverse primer, 25 nM of the M13 tagged forward primer and 100 nM of the dye labeled M13 forward primer. The cycling parameters were an initial denaturation step for 1 min at 94 C, 33 cycles of denaturation at 94 C for 40 s, annealing at 58 C for 40 s, extension at 72 C for 20 s, followed by a final synthesis step at 72 C for 30 min. PCR products were pooled based on the fluorescent phosphoramidite dyes used (VIC, NED, PET, FAM) to a 1:10 dilution. The pooled PCR products were added to 9.9 μL of Hi-Di formamide and 0.1 μL of GeneScan™ 500 LIZ™ size standard (Life Technologies). Samples were denatured at 94 C for 5 min. Microsatellite fragments were analyzed on an ABI 3730 DNA Analyzer. Analysis was performed using Peak Scanner Software v1.0 (Applied Biosystems, Foster City, California). The 32 markers were successfully scored in the crosses except for two markers, Fe176_AGG12-F and Fe179_CTT6-F, due to mismatches in the parental strains NFVe47 and FRT5LL7-Albino. Each progeny colony was also tested for mating type as described above using a multiplex reaction including the housekeeping gene *TUB2* as previously described (Young et al. 2018).

The proportion of MAT1-1-1 and MAT1-2-1 isolates was tested for equilibrium (the null hypothesis being 1:1 ratio) using a χ2 test to determine whether the proportions departed significantly from a 1:1 ratio (Everitt 1992). Due to the limitations of the χ2 test when analyzing small sample sizes (generally populations with expected values <5), an exact binomial test (two-tailed) for goodness-of-fit was used to determine whether observed mating type frequencies within populations deviated from the 1:1 ratio. Analyses of mating type data were performed using SAS V9.4 (SAS Institute, Cary, NC).

### Confirmation of the causative mutation for the albino phenotype

Progeny from the crosses with the FRT5LL7-Albino parent were evaluated for the single nucleotide insertion in the melanin biosynthesis gene *PKS1* encoding a polyketide synthetase, the causative mutation of the albino phenotype (Winter et al. 2019). DNA from each of the parents and the progeny were amplified with the primers alm-F (5’- CAGTGTGATGGCCTTGACTATCAG-3’) and alm-R (5’-TTCTCCAACAAGTCCCAG-3’) that results in a 583 bp wild-type fragment spanning the proposed causative mutation. The PCR products were cleaned up using QIAquick PCR Purification Kit (QIAGEN Inc.) according to manufacturer’s instructions and then direct sequenced using Sanger sequencing technology in the Genomics Core Facility, Noble Research Institute, LLC.

## RESULTS

### Mating assay

Mating type for six isolates were previously identified (Young et al. 2018) and the remaining 10 isolates were characterized as *MAT1-1* or *MAT1-2* based on multiplex PCR (TABLE 1). Different isolates representing *MAT1-1* and *MAT1-2* types were paired (TABLE 2) and observed for production of pseudothecia. Fruiting bodies were not observed on PDA. Immature fruiting bodies were observed after 6 months incubation (2 mo at 24 C and 4 mo at 4 C) during the preliminary experiments on oatmeal agar (Fig. 2). Once transferred to 24 C with a 12 h photoperiod, maturation of the pseudothecia occurred, and mature asci and ascospores were observed (Fig. 3). The squash mounts of the pseudothecia confirmed maturation by the presence of fully developed asci and ascospores. Pseudothecia were observed between all paired isolates of opposite mating types. Pseudothecia never formed between self-pairings or non-self-same mating type pairings. Pseudothecia formed on the bottom, top and embedded throughout the agar between isolates of opposite mating types, regardless as to the orientation of the Petri dish during incubation. Depending on the isolate, some pseudothecia would form closer to one parent over the other, which may indicate differences in fertility between isolates.

**FIGURE 2.**
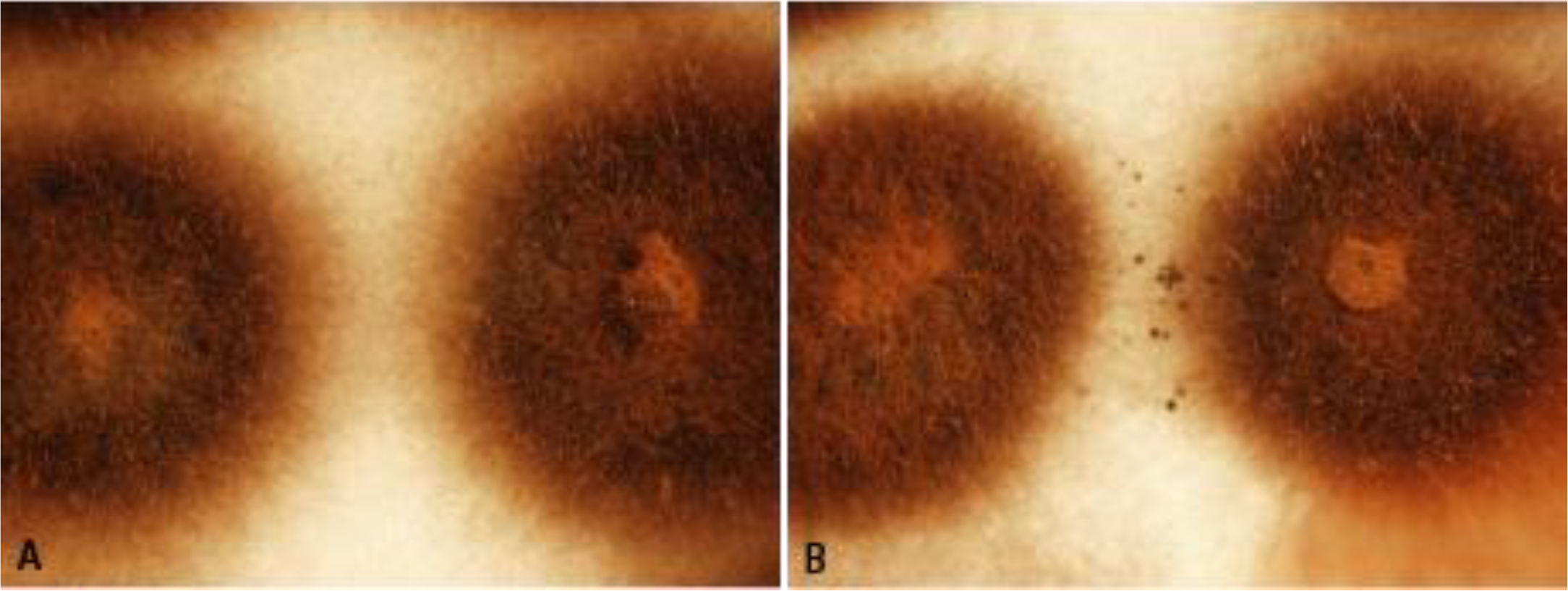
Zone observed between pairings after 4 mo cold treatment (4 C). **A**) Same mating types (ALB1L5 × ALB1L5) and **B**) opposite mating types (ALB1L5 × ALB2L1).

**FIGURE 3.**
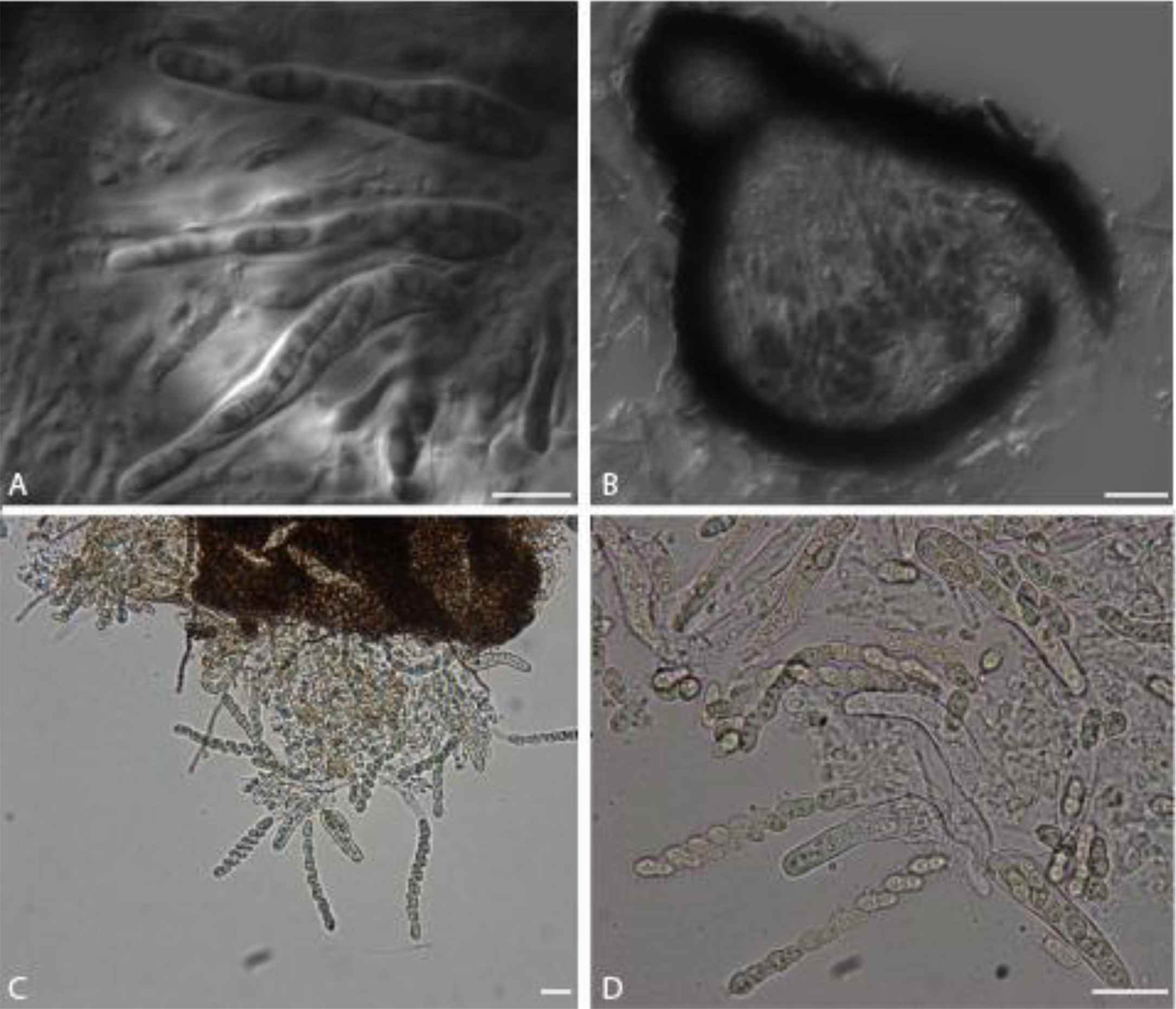
Sexual reproductive structures in *Venturia effusa*. **A**) DIC image of a section of a pseudothecium from a cross between NFVe23 × NFVe24 3wk at 24C 12h photoperiod post cold incubation with ascospores in asci. **B**) DIC image of a section of a pseudothecium from NFVe23 × NFVe49 5wk at 24C 12h photoperiod post cold incubation. Scale bars: A = 10 μm, B = 20 μm. **C** and **D**) Squash mount of pseudothecium from NFVe23 × NFVe49 6 wk at 24 C following 4 C incubation. Numerous asci about to release ascospores. Scale bars = 20 μm.

The morphology of the pseudothecia, asci and ascospores observed were consistent with descriptions of the genus *Venturia* (Von Arx 1952; Zhang et al. 2011) (Fig. 3). Orientation of the ascospores in each ascus are similar to *V. pyrina* rather than *V. inaequalis* with the larger end of the spore facing towards the ostiole. Ascospores of *V. effusa*, like *V. pyrina,* are septate in the lower third in contrast to *V. inaequalis* ascospores which are septate in the upper third (Barr 1968). Pseudothecia from *V. inaequalis* have been described as being with or without setae (MacHardy 1996) and may have setae surrounding the ostiole according to Gadoury and MacHardy (1985). We observed pseudothecia with and without setae (Fig. 4A–D).

**FIGURE 4.**
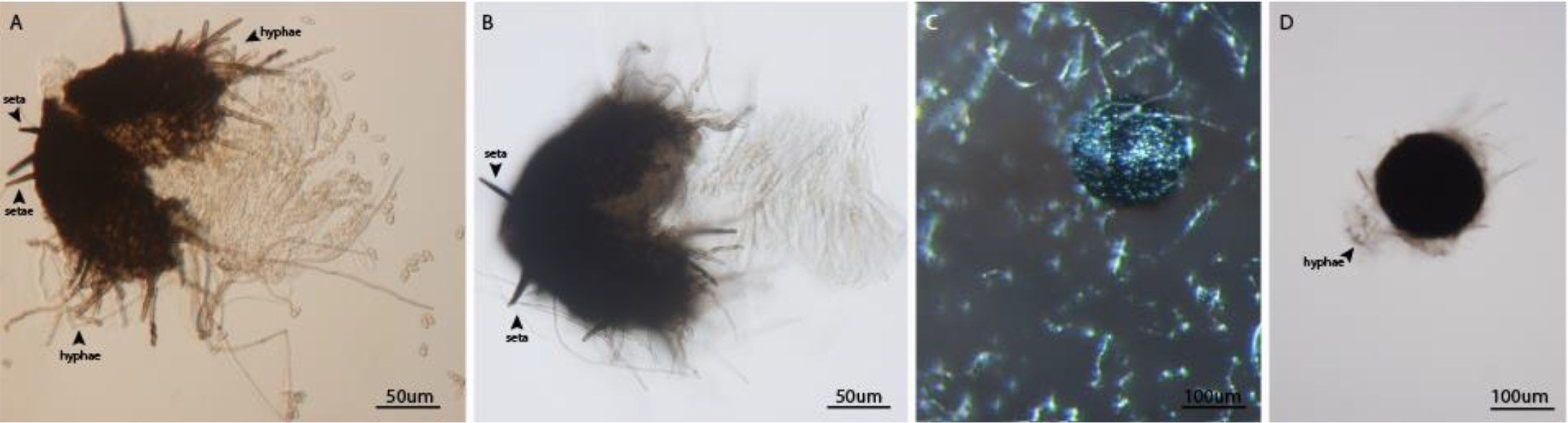
Pseudothecia of *Venturia effusa* with and without setae. Pseudothecia developed in the oatmeal agar and therefore hyphae were typically present on all pseudothecia, whereas setae can be distinguished from hyphae as highly pigmented bristle like hairs. **A** and **B**) Examples of setae observed on pseudothecia. Scale bars = 50 μm. **C** and **D**) Examples of pseudothecia without setae. Scale bars = 100 μm.

### Effect of time on pseudothecia development

Observations were made to assess the development of pseudothecia and the maturation of asci after varying the length of the cold treatment. The pairings observed after one month of cold treatment showed no signs of fruiting bodies or fruiting body initials. The pairings observed after two months of cold treatment had both pseudothecia and smaller ‘hyphal knots’, which may be protopseudothecia. The pseudothecia appeared to remain immature or partially mature, with few ascospores observed. All pairings observed after 4 mo had some asci with ascospores. The pseudothecia became more mature (increase in number of asci with ascospores) after a minimum 2 wk incubation up to 6 wk at 24 C.

### Multilocus genotyping of ascospore progeny

Progeny from independent sexual crosses were evaluated to determine if sexual reproduction produced recombinant progeny (TABLE 3). A total of 32, 13 and 30 single ascospore progeny were isolated from three crosses between isolates NFVe23 (cv. Pawnee) × NFVe24 (native), FRT5LL7-Albino (cv. Desirable) × NFVe47 (cv. Choctaw) and FRT5LL7-Albino (cv. Desirable) × NFVe78 (cv. Pawnee), respectively. The percentage of unique multilocus genotypes for NFVe23 × NFve24, FRT5LL7-Albino × NFVe47 and FRT5LL7-Albino × NFVe78 were 88% (4 possible clones), 100% and 93% (2 possible clones), respectively. The mating type ratios (*MAT1-1*: *MAT1-2*) were 17:15, 9:4 and 14:16. Mating types of all three crosses were in gametic equilibrium based on both the χ2 test and the exact binomial test (TABLE 4). The phenotype ratio was 5:8 and 10:20 (albino:wild type), which although in statistical equilibrium (data not shown), may indicate some numerical imbalance in frequency which could be due to wild type spores being preferentially selected over albino spores, perhaps due to difficulty observing hyaline, albino spores on the media.

**Table 3.** Genotyping ascospore progeny produced from *in vitro* sexual crosses using 32 microsatellite markers, mating type and phenotype if applicable.

| Cross | No. of progeny | No. of markers screened | No. of successful markers | No. of informative markers | Albino : WT | MAT1-1: MAT1-2 | Unique multilocus genotypes based on microsatellite | Unique multilocus genotypes based on microsatellite/ MAT/ phenotype |
| --- | --- | --- | --- | --- | --- | --- | --- | --- |
| NFVe 23 (Pawnee) × NFVe 24 (Native) | 32 | 32 | 32 | 19 | N/A | 17:15 | 25 | 28 |
| Albino (Desirable) × NFVe 47 (Choctaw) | 13 | 32 | 30 | 25 | 5:8 | 9:4 | 13 | 13 |
| Albino (Desirable) × NFVe 78 (Pawnee) | 30 | 32 | 31 | 20 | 10:20 | 14:16 | 27 | 28 |

**Table 4.** The mating type frequencies of the different crosses of isolates of *Venturia effusa.*

| Cross | *MAT1-1-1: MAT1-2-1* | | $\chi^2$ (P)<sup>a</sup> | p<sup>b</sup> |
| --- | --- | --- | --- | --- |
|  | Number | % |  |  |
| NFVe23 (cv. Pawnee) × NFVe24 (native) | 17:15 | 53.1:46.9 | 0.1 (0.7) | 0.7 |
| FRT5LL7-Albino (cv. Desirable) × NFVe47 (cv. Choctaw) | 9:4 | 69.2:30.8 | 1.9 (0.2) | 0.2 |
| FRT5LL7-Albino (cv. Desirable) × NFVe78 (cv. Pawnee) | 14:16 | 46.7:53.3 | 0.1 (0.7) | 0.7 |

<sup>a</sup> $\chi^2$ value based on a contingency $\chi^2$ between clone-corrected mating type ratios with 1 degree of freedom. The $\chi^2$ value is based on a 1:1 ratio with 1 degree of freedom; the P-value is in parentheses.
<sup>b</sup> Probability from an exact binomial analysis (two-tailed) to test whether mating type frequencies deviate significantly from a 1:1 ratio (more appropriate for comparing small populations than a $\chi^2$ test).

Genes involved in melanin biosynthesis were identified from the *Venturia effusa* genome sequence of 3Des10b (Bock et al. 2016b) using TBlastN with orthologs from the *Alternaria alternata* melanin gene cluster (Kimura and Tsuge 1993; Fetzner et al. 2014). Three genes, *PKS1* encoding a polyketide synthetase with 79% similarity to ALM1 (PKSA), *CRM1* encoding a C2H2 Zn^2+^ finger transcription factor with 71% similarity to CRMA, and *BRM2* encoding a T3HN reductase with 85% similarity to BRM2 were located in a gene cluster in a similar orientation to *A. alternata*. Evaluation of the FRT5LL7-Albino genome sequence revealed the melanin gene cluster was located on chromosome 6. Comparison of each gene to the 3Des10b sequence revealed a critical mutation in *PKS1* from FRT5LL7-Albino as a single base insertion (Winter et al. 2019) (Fig. 5). All progeny with the albino phenotype (n=15) have the identical mutation found in the FRT5LL7-Albino *PKS1,* confirming this mutation is the likely cause of the albino phenotype.

**FIGURE 5.**
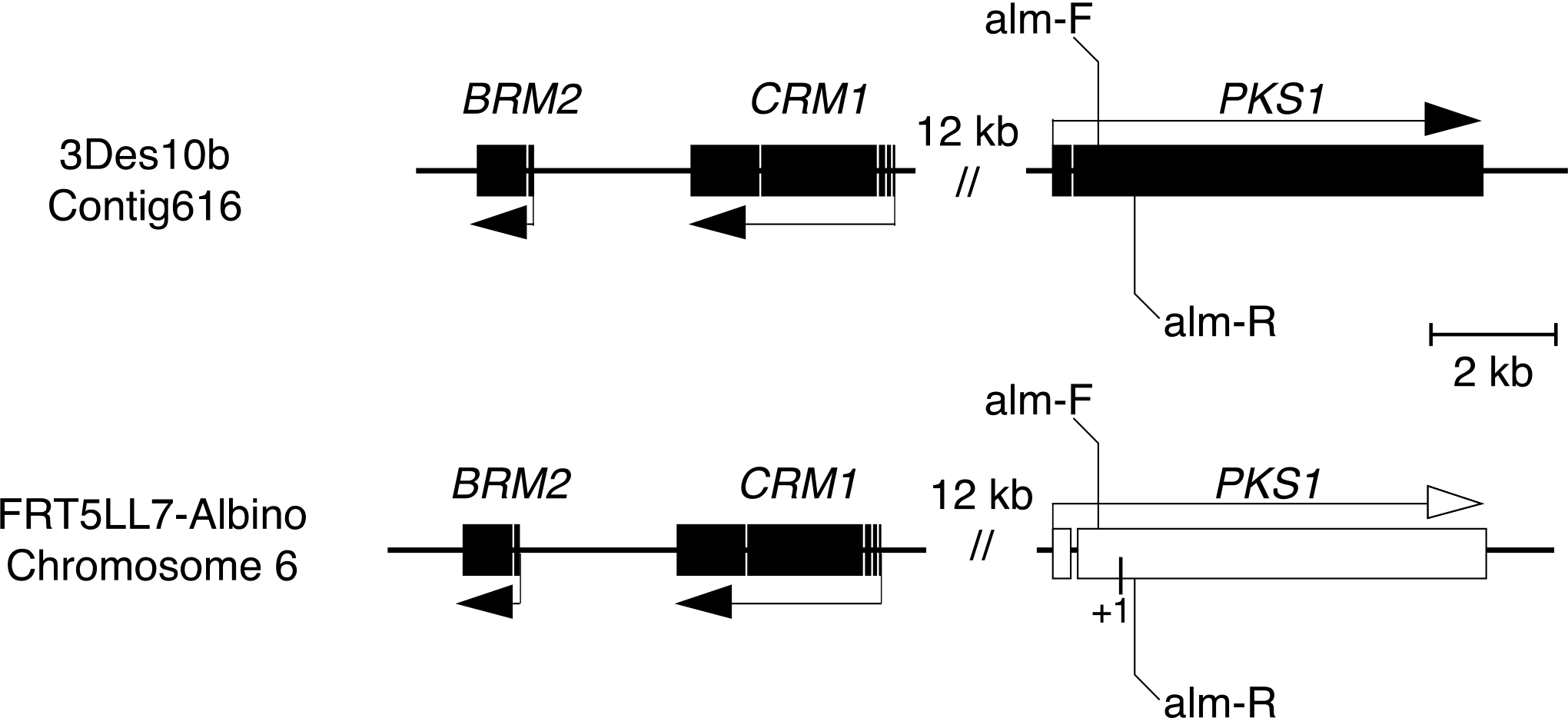
Genome organization of melanin biosynthesis genes. Coding regions are represented as black boxes on the sequence (black line). Genes are represented as an arrow pointing in the direction of transcription. The approximate location of the mutation in FRT5LL7-Albino *PKS1* gene is represented by +1, indicating the insertion of a single nucleotide, which resulted in a frameshift that rendered the *PKS1* gene nonfunctional. Primers used in a PCR screen to test progeny for the albino genotype are alm-F and alm-R. *CRM1* and *PKS1* are the equivalent of *CRMA* and *ALM1* or *PKSA* in *Alternaria alternata*, respectively.

## DISCUSSION

This is the first report of successful mating of *V. effusa in vitro* through direct contact of isolates of opposite mating types on OMA. We demonstrated production of pseudothecia and mature, viable ascospores between isolates from different locations and pecan cultivars, which was reproducible (5 crosses between isolates from the same cultivars, 5 crosses between isolates from different cultivars). *V. effusa* was able to complete the sexual cycle after a minimum 2 mo cold treatment, but with a greater number of asci maturing after a 4 mo cold treatment. Maturation of asci and ascospores did require an increase in temperature for a minimum of 2 wk after the cold period, with an increase in the number of mature asci as the incubation time at 24 C increased (observed up to 6 wk at 24 C).

Ascospore viability was confirmed and genotyping the progeny demonstrated genetic recombination through sexual reproduction. Evidence that *V. effusa* can reproduce sexually supports the population studies that have identified moderate to high genetic diversity along with mating type ratios in equilibrium (Bock et al. 2017b; Bock et al. 2018; Young et al. 2018). The progeny from the crosses revealed unique multilocus genotypes based on the microsatellite analysis where the location of the microsatellite markers are known to be distributed across multiple chromosomes (Winter et al. 2019). These observations support (and provides a basis for) the results from studies of field populations of *V. effusa* showing characteristics of population genetic diversity and structure suggestive of sexual reproduction. The recombinant genotypes resulting from sexual reproduction could lead to a more adaptable fungal population (McDonald and Linde 2002; McDonald and Mundt 2016).

Ascospores of *Venturia* species can play an important role not just as a vehicle of novel diversity, but also as a component of the epidemiology of the disease (González-Domínguez et al. 2017). Studies from other *Venturia* species may aid in identifying the role of ascospores in the epidemiology of pecan scab. For example, conidia of *Venturia nashicola* have been considered the primary inoculum for pear scab, but recent studies suggest that ascospores may play a more important role in the epidemiology of this disease. Researchers have focused their efforts on studying pseudothecia formation and ascospore discharge to better understand the role of ascospores. Results showed that pseudothecia formation in pear scab in northern China was influenced by rain events in either winter or early spring with ascospore release occurring by late May (Lian et al. 2006). Further studies by (Lian et al. 2007) found that free water or 100% RH was required for ascospore release with approximately 80% of ascospores being released within the first hour after being wetted.

Asscopores of *V. inaequalis* play a significant role in epidemics of apple scab (MacHardy 1996), and some control measures can effectively target the sexual stage (Holb 2006). *In vitro* sexual reproduction in *V. inaequalis* may have similar requirements to that identified for *V. effusa*, but these findings have not been effectively reproduced. Early studies showed pseudothecia of *V. inaequalis* would form in culture on OMA when stored at 10 C for 5 months (Jones 1914a; Jones 1914b). However, others have been unable to produce ascospores *in vitro* without using apple leaves, and have only seen pseudothecia initials on oatmeal agar (Rudloff 1934; Schmidt 1935; Herbst et al. 1937). Infected leaves have been used under controlled growth conditions to determine environmental requirements. Moisture and temperature were factors in pseudothecial development of *V. inaequalis* on apple in North Carolina (James and Sutton 1982). In addition, moisture and temperature from Feb to Apr influenced the maturation of pseudothecia with the optimum temperature range for ascogonial development of 8-12 C and 16-18 C for ascospore maturation (James and Sutton 1982). Earlier studies found that pseudothecia production was greatest at 4 C and asci without ascospores at 15 C (Ross and Hamlin 1962). With the identification of a sexual stage in *V. effusa*, disease management may now align more closely to management practices for apple scab.

If the *V. effusa* life cycle is similar to those of *V. inaequalis* and *V. nashicola*, one would suspect the source of overwintering pseudothecia would be on dead pecan leaves. An ascomycetous fungus has been observed on dead pecan leaves, but the fungus was not identified (Demaree 1924). In Oklahoma, we have observed initial infections on the abaxial side of leaves lowest in the canopy (approximately 2 m) in early May to early June, which leads us to hypothesize that the inoculum source may in fact be on the orchard floor. Demaree (1924) did note an evident reduction in scab after a fire moved through a heavily infected orchard, as well as control of the disease in plowed orchards where the leaf, twig and shuck debris were buried under the soil compared to unplowed orchards. We have identified an ascomycetous fungus, *Mycosphaerella*, in the pecan leaf litter (Charlton and Young, Unpublished), which has introduced an additional challenge to identifying the role of *V. effusa* ascospores in the disease cycle of pecan scab. *Mycosphaerella carygena*, cause of downy spot, and *V. effusa* appear to have morphologically similar pseudothecia and ascospores. The timing of *M. carygena* ascospore release has been established as early spring prior to budbreak (early April) (Goff et al. 1987), around the same time we would expect *V. effusa* ascospores to be released. The morphological characteristics of the culture *in vitro* can be used to distinguish *V. effusa* and *M. carygena*, but can take up to one week. Due to the similarity of the ascospores, it will likely be challenging to identify the role of the sexual spores in the epidemiology of pecan scab.

The discovery of the sexual stage of *V. effusa* could have profound implications on management strategies to control pecan scab. Additional studies are required to determine the optimal temperature ranges for pseudothecial development and asci maturation in *V. effusa in vitro*. Future studies that focus on the time and environmental factors required to initiate and develop mature pseudothecia could provide more definitive requirements for the sexual cycle and ascospore release under field conditions. Identification of where *V. effusa* overwinters and the location of pseudothecia development is critical to develop management strategies to reduce the primary inoculum in the disease cycle, as has been described for *V. inaequalis*. With further knowledge of the sexual stage of *V. effusa* in the field, we have an opportunity to develop additional science-based management tools to control scab in pecan.

## ACKNOWLEDGEMENTS

We thank Jin Nakashima and the Cellular Imaging Core Facility, Noble Research Institute LLC for technical support and Rachael Davis, Noble Research Institute LLC for graphic design. The authors also wish to thank Basil Savage for allowing access to his orchard to make observations and collect samples throughout the year as well as Dr. Charles Rohla, Noble Research Institute LLC, for helping collect samples. CHB and MZ were supported by the USDA-ARS through CRIS project 6606-21220-011–00D.

This article reports the results of research only. Mention of a trademark or proprietary product is solely for the purpose of providing specific information and does not constitute a guarantee or warranty of the product by the U.S. Department of Agriculture and does not imply its approval to the exclusion of other products that may also be suitable.

## LITERATURE CITED

Barr ME. 1968. The Venturiaceae in North America. Canadian Journal of Botany 46:799–864.

Bock C, Chen C, Yu F, Stevenson K, Arias R, Wood B. 2016a. Characterization of microsatellites in *Fusicladium effusum*, cause of pecan scab. Forest Pathology 46:600–609.

Bock C, Brenneman T, Wood B, Stevenson K. 2017a. Challenges of managing disease in tall orchard trees-pecan scab, a case study. CAB Reviews 12:1–18.

Bock CH, Chen C, Yu F, Stevenson KL, Wood BW. 2016b. Draft genome sequence of *Fusicladium effusum*, cause of pecan scab. Standards in genomic sciences 11:36.

Bock CH, Hotchkiss MW, Young CA, Charlton ND, Chakradhar M, Stevenson KL, Wood BW. 2017b. Population genetic structure of *Venturia effusa*, cause of pecan scab, in the southeastern United States. Phytopathology 107:607–619.

Bock CH, Young CA, Stevenson KL, Charlton ND. 2018. Fine-scale population genetic structure and within-tree distribution of mating types of *Venturia effusa*, cause of pecan scab in the United States. Phytopathology 108:1326–1336.

Brock J and Brenneman TB. 2014. Pecan. Page 14 in: UGA Ext. Annu.Publ. 102–7, University of Georgia.

Demaree JB. 1924. Pecan scab with special reference to sources of the early spring infections. Journal of Agricultural Research 28:321–333.

Everitt BS. 1992. The analysis of contingency tables. London: Chapman and Hall/CRC.

Fetzner R, Seither K, Wenderoth M, Herr A, Fischer R. 2014. *Alternaria alternata* transcription factor CmrA controls melanization and spore development. Microbiology 160:1845–1854.

Fisher EE. 1961. *Venturia carpophila* sp. nov., the ascigerous state of the apricot freckle fungus. Transactions of the British Mycological Society 44:337–342.

Gadoury DM, MacHardy WE. 1985. Negative geotropism in *Venturia inaequalis*. Phytopathology 75:856–859.

Gladieux P, Zhang X-G, Afoufa-Bastien D, Sanhueza R-MV, Sbaghi M, Le Cam B. 2008. On the origin and spread of the scab disease of apple: out of central Asia. PloS one 3:e1455.

Gladieux P, Zhang XG, Róldan‐Ruiz I, Caffier V, Leroy T, Devaux M, Van Glabeke S, Coart E, Le Cam B. 2010. Evolution of the population structure of *Venturia inaequalis*, the apple scab fungus, associated with the domestication of its host. Molecular Ecology 19:658–674.

Goff W, Drye C, Miller R. 1987. Ecology and epidemiology of pecan downy spot. Phytopathology 77:491–496.

González-Domínguez E, Armengol J, Rossi V. 2017. Biology and epidemiology of *Venturia* species affecting fruit crops: a review. Frontiers in plant science 8:doi.org/10.3389/fpls.2017.01496.

Gottwald T. 1982. Spore discharge by the pecan scab pathogen, *Cladosporium caryigenum*. Phytopathology 72:1193–1197.

Gottwald T, Bertrand P. 1982. Patterns of diurnal and seasonal airborne spore concentrations of *Fusicladium effusum* and its impact on a pecan scab epidemic. Phytopathology 72:330–335.

Gottwald T, Bertrand P. 1983. Effect of time of inoculation with *Cladosporium caryigenum* on pecan scab development and nut quality. Phytopathology 73:714–718.

Gottwald T. 1985. Influence of temperature, leaf wetness period, leaf age, and spore concentration on infection of pecan leaves by conidia of *Cladosporium caryigenum*. Phytopathology 75:190–194.

Herbst W, Rudloff C, Schmidt M. 1937. Vergleichend-morphologische Studien an verschiedenen Venturiaarten. Die Gartenbauwissenschaft 11:183–207.

Holb IJ. 2006. Effect of six sanitation treatments on leaf litter density, ascospore production of *Venturia inaequalis* and scab incidence in integrated and organic apple orchards. European Journal of Plant Pathology 115:293–307.

James J, Sutton T. 1982. Environmental factors influencing pseudothecial development and ascospore maturation of *Venturia inaequalis*. Phytopathology 72:1073–1080.

Jones F. 1914a. Perithecia in cultures of *Venturia inaequalis*. Phytopathology 4:52–53.

Jones FR. 1914b. Is the *Venturia* on the Wisconsin wild crab, *Pyrus coronaria*, identical with the *Venturia inequalis* of the cultivated apple. University of Wisconsin: University of Wisconsin, Madison.

Kimura N, Tsuge T. 1993. Gene cluster involved in melanin biosynthesis of the filamentous fungus *Alternaria alternata*. Journal of Bacteriology 175:4427–4435.

Lian S, Li BH, Xu XM. 2006. Formation and development of pseudothecia of *Venturia nashicola*. Journal of Phytopathology 154:119–124.

Lian S, Li BH, Dong XL, Li BD, Xu XM. 2007. Effects of environmental factors on discharge and germination of ascospores of *Venturia nashicola*. Plant Pathology 56:402–411.

Linde CC, Zhan J, McDonald BA. 2002. Population structure of *Mycosphaerella graminicola*: from lesions to continents. Phytopathology 92:946–955.

Littrell R. 1976. Resistant pecan scab strains to benlate and pecan fungicide management. Pecan South 3:335–337.

MacHardy WE. 1996. Apple scab: biology, epidemiology, and management. St. Paul, Minnesota: American Phytopathological Society (APS Press). p. 545.

McDonald BA, McDermott JM. 1993. Population genetics of plant pathogenic fungi. Bioscience 43:311–319.

McDonald BA, Pettway RE, Chen R, Boeger J, Martinez J. 1995. The population genetics of *Septoria tritici* (teleomorph *Mycosphaerella graminicola*). Canadian Journal of Botany 73:292–301.

McDonald BA, Linde C. 2002. Pathogen population genetics, evolutionary potential, and durable resistance. Annual Review of Phytopathology 40:349–379.

McDonald BA, Mundt CC. 2016. How knowledge of pathogen population biology informs management of Septoria tritici blotch. Phytopathology 106:948–955.

Milgroom MG. 1996. Recombination and the multilocus structure of fungal populations. Annual Review of Phytopathology 34:457–477.

Passey T, Robinson J, Shaw M, Xu XM. 2017. The relative importance of conidia and ascospores as primary inoculum of *Venturia inaequalis* in a southeast England orchard. Plant Pathology 66:1445–1451.

Reynolds KL, Brenneman TB, Bertrand PF. 1997. Sensitivity of *Cladosporium caryigenum* to propiconazole and fenbuconazole. Plant Disease 81:163–166.

Ross R, Hamlin S. 1962. Production of perithecia of *Venturia inaequalis* (Cke.) Wint. on sterile apple leaf discs. Canadian Journal of Botany 40:629–635.

Rossman AY, Allen WC, Castlebury LA. 2016. New combinations of plant-associated fungi resulting from the change to one name for fungi. IMA fungus 7:1–7.

Rudloff C. 1934. *Venturia inaequalis* (Cooke) Aderhold. III. Zur Formenmannigfaltigkeit des Pilzes. Die Gartenbauwissenschaft 9:105–119.

Schmidt M. 1935. *Venturia inaequalis* (Cooke) Aderhold. IV. Weitere Beiträge zur Rassenfrage beim Erreger des Apfelschorfes. Die Gartenbauwissenschaft 9:364–389.

Schubert K, Ritschel A, Braun U. 2003. A monograph of *Fusicladium* s. lat. (Hyphomycetes). Schlechtendalia 9:1–132.

Standish J, Avenot H, Brenneman T, Stevenson K. 2016. Location of an intron in the cytochrome b gene indicates reduced risk of QoI fungicide resistance in *Fusicladium effusum*. Plant Disease 100:2294–2298.

Standish J, Brenneman T, Stevenson K. 2018. Dynamics of fungicide sensitivity in *Venturia effusa* and fungicide efficacy under field conditions. Plant Disease 102:1606–1611.

Stevenson K, Bertrand P. 2001. Within-season dynamics of yield loss due to pecan scab fruit infections. Phytopathology 91:S85.

Umemoto S. 1990. Infection sources in Japanese pear scab (*Venturia nashicola*) and their significance in the primary infection. Japanese Journal of Phytopathology 56:658–664.

Von Arx J. 1952. Studies on *Venturia* and related genera. Tijdschrift over Plantenziekten 58:260–266.

Wells L. 2017. Pecan: America’s Native Nut Tree. University of Alabama Press.

Winter DJ, Charlton ND, Krom N, Shiller J, Bock CH, Cox MP, Young CA. 2019. Chromosome-level genome reference of *Venturia effusa*, causative agent of pecan scab. bioRxiv 746198 doi: https://doi.org/10.1101/746198

Winter G. 1885. New North American fungi. The Journal of Mycology 1:101–102.

Wood BW, Payne JA, Grauke LJ. 1990. The rise of the US pecan industry. HortScience 25:594–723.

Young CA, Bock CH, Charlton ND, Mattupalli C, Krom ND, Bowen JK, Templeton MD, Plummer K, Wood BW. 2018. Evidence for sexual reproduction: identification, frequency and spatial distribution of *Venturia effusa* (pecan scab) mating type idiomorphs. Phytopathology 108:837–946.

Zhang Y, Crous PW, Schoch CL, Bahkali AH, Guo LD, Hyde KD. 2011. A molecular, morphological and ecological re-appraisal of *Venturiales*―a new order of *Dothideomycetes*. Fungal Diversity 51:249–277.

